# Development of a High-Throughput Screening Method for Anti-NNV Drugs

**DOI:** 10.1101/2025.09.30.679469

**Authors:** Mingzhu Liu, Jingu Shi, Yu Gao, Weijiang Xu, Lin Huang, Huapu Chen, Yong Zhang, Jia Cai, Xiaowen Zhu, Shuyu Han, Jinkun Xie, Qing Yu, Pengfei Li

## Abstract

Nervous necrosis virus (NNV) is an infectious pathogen, characterized by rapid infection and high mortality, commonly encountered in the aquaculture industry. Therefore, there is an urgent need to develop effective antiviral drugs against NNV, which is crucial for targeted treatment strategies, pathogen control, and loss reduction. Screening of effective antiviral active compounds, namely drug precursors, is key to developing highly efficient drugs against NNV. Reverse transcription-quantitative real-time PCR (RT-qPCR) is widely applied to screen and evaluate various substances to prevent and control virus infection. However, RT-qPCR is a cumbersome procedure and not suitable for high-throughput rapid screening. The aptamer TNA1c, which specifically binds to NNV-infected cells, was used to construct a target-driven activatable aptamer probe (TAA). Then, the TAA probe was applied to establish a high-throughput screening (TAA-HTS) method for efficient evaluation of substances against NNV infection. TAA-HTS technology achieved rapid, sensitive, and specific screening of bioactive substances with significant anti-NNV effects. As compared to commonly used analytical methods, such as RT-qPCR, TAA-HTS has the advantages of easy operation and high sensitivity and specificity. The findings of this study provide data support and a theoretical basis for the development of effective antivirus preparations.

**IMPORTANCE:** In this study, a target-driven activatable aptamer probe was employed to establish a high-throughput screening method for anti-NNV drugs. This method enables rapid, sensitive, and specific screening of anti-NNV compounds while reducing background interference. Although RT-qPCR is currently the most widely used technique for antiviral drug screening and validation, the screening approach developed in this study significantly shortens screening time while achieving results consistent with those of RT-qPCR. This demonstrates that the screening method presented here has the potential to become a universal strategy for future drug screening against viral diseases. Moreover, this study not only provides a novel technical solution for NNV control but also demonstrates the potential of aptamer-based probes in high-throughput antiviral screening, serving as a model for combating viral pathogens in aquaculture and other fields.

---

*Epinephelus* is a genus of economically important mariculture fish species native to tropical and subtropical seas and oceans. With the rapid and intensive development of the aquaculture industry, grouper production has continued to increase in recent years. However, farmed groupers are prone to various diseases, especially infection by the nervous necrosis virus (NNV), which has a fatality rate as high as 99%, resulting in huge economic losses to the aquaculture industry(1, 2). NNV is a member of the Nodaviridae family of nonenveloped positive-strand RNA viruses with diameters of 25–30 nm. After infection, NNV can cause damage to the nervous system of fish, causing dysmotility and abnormal swimming, which seriously affect growth and survival. Therefore, it is particularly important to develop effective and safe antiviral drugs to maintain the health and sustainable development of farmed groupers.

Herbs are important components of traditional Chinese medicine (TCM) that contain a diverse array of chemical constituents, including fatty acids, amino acids, carbohydrates, terpenoids, flavonoids, and alcohols, which exhibit broad biological activities, especially antiviral effects(3). Active compounds of TCM, such as curcumin, dimethoxy curcumin, chlorogenic acid, caffeic acid, luteolin, and quercetin, have demonstrated inhibitory effects against the Singapore grouper iridovirus (SGIV) (4-6). Additionally, epigallocatechin gallate has been shown to inhibit replication of the grass carp reovirus and largemouth bass iridovirus (7), while luteolin is reportedly effective against white spot syndrome (8) and quercetin can induce expression of interferon-related genes and enhance host immune responses (9, 10). These bioactive compounds combat viral infections by various mechanisms to promote the healthy growth of aquatic animals, primarily by inhibition of viral adsorption and entry, interference with viral replication, and enhancement of host immune responses. The application of the bioactive compounds of TCM contributes to the sustainability of farmed fish with minimal side effects. Therefore, various herbs used in TCM have been screened as potential antiviral agents for use in the aquaculture industry.

Currently, quantitative real-time fluorescent polymerase chain reaction (RT-qPCR) is predominantly utilized for screening of antiviral drugs. Although highly accurate and reproducible, RT-qPCR requires complex procedures and lengthy processing times. Recent studies have shown that aptamers possess the ability to specifically recognize target substances. Consequently, many researchers have used aptamers for the detection of aquaculture pathogens. For instance, the aptamer LYGV1 can specifically recognize SGIV(11), while A5 and GBN34 can specifically identify NNV (12, 13). Additionally, LA38s and LA13s are capable of specifically recognizing the largemouth bass iridovirus (14). At present, most aptamers are labeled with fluorophores at one end, thus remaining in an activated state (emitting light), necessitating multiple washing steps to minimize the impact of unbound probes, which can result in high background signals. In this study, we employed a detection method by labeling both ends of the probe with a fluorophore and a quencher group, thereby maintaining the probe in a non-emissive state (quiescent) at the outset. Then, conformational alterations triggered by aptamer recognition of the target leads to signal alteration as an “activable probe detection method” (15-17). This method reduces the washing steps, while shortening the detection time and minimizing the influence of background signals.

The aim of this study was to assess the applicability of a high-throughput screening method based on aptamers (TAA-HTS) to identify drugs effective against NNV. An active aptamer probe was employed to facilitate rapid screening of bioactive compounds to inhibit replication of NNV. This technology is expected to accelerate the development of antiviral drugs, enhance screening efficiency, and reduce development costs, thereby providing effective preventive and therapeutic measures for the healthy farming of groupers.

**FIG 1.**
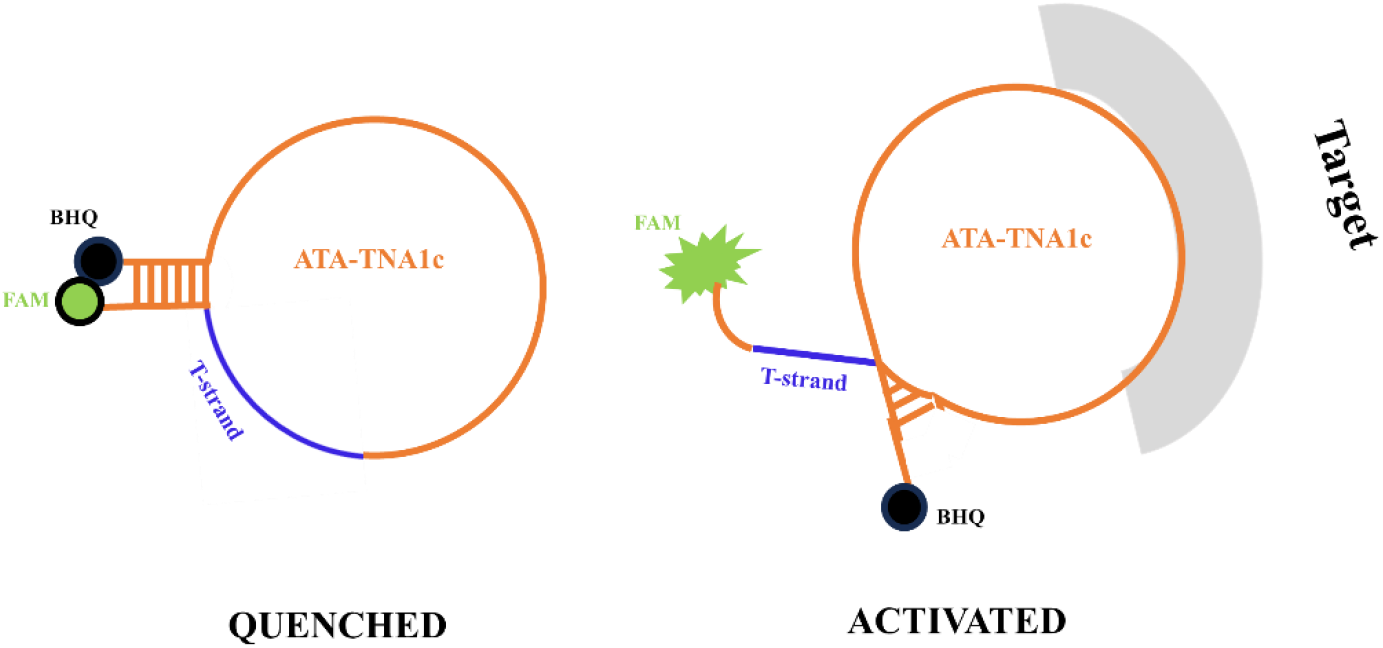
Schematic representation of the novel aptamer probe TAA-TNA1c.

## RESULTS

### Specificity analysis of TAA-TNA1c for NNV infection

As shown in FIG 2, TAA-TNA1c specifically recognized cells infected with NNV, but not SGIV or the uninfected control cells. These results indicate that TAA-TNA1c is highly specific for NNV.

**FIG 2.**
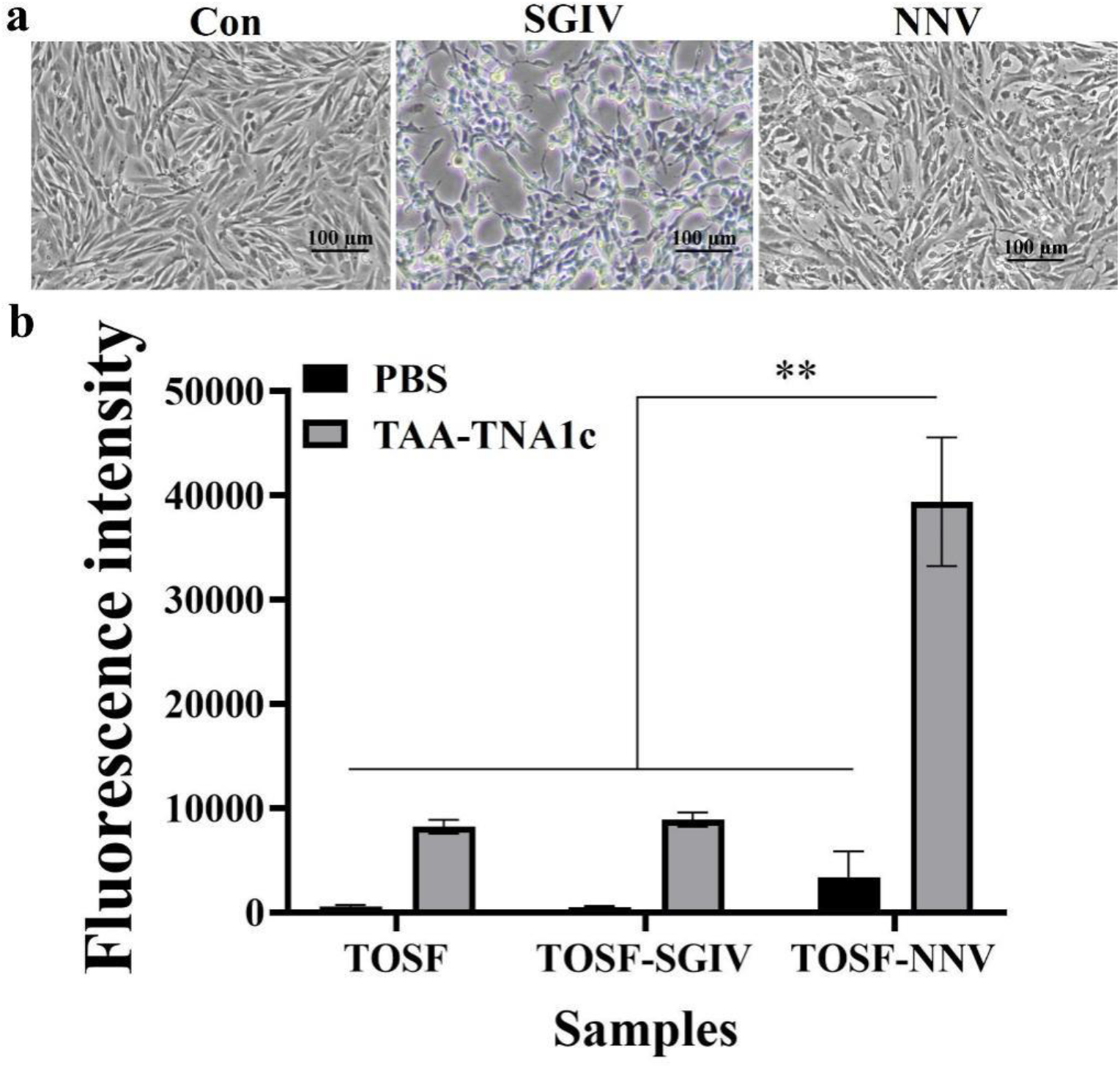
Specificity analysis of TAA-TNA1c. a. Morphological changes in TOSP infected with NNV or SGIV were assessed by light microscopy. b. The efficacy of TAA-TNA1c in detecting NNV infection was evaluated using a fluorescent microplate reader.

### Effects of different concentrations of NNV

Fluorescence generated by the TAA-TNA1c probe at varying concentrations of NNV was measured with a fluorescence microplate reader (FIG 3a). At probe concentrations of 0-12µL/mL, the fluorescence of the probe-treated groups increased progressively with increasing NNV concentrations. Notably, the fluorescence values were significantly higher in the TOSP-NNV group with the probe than without. In contrast, there were no significant differences in fluorescence between the probe-treated and untreated groups of TOSP cells. CP gene expression levels at different NNV concentrations are shown in FIG 3b. The results demonstrate that CP expression levels increased with higher NNV concentrations, showing significant differences as compared to the control group.

**FIG 3.**
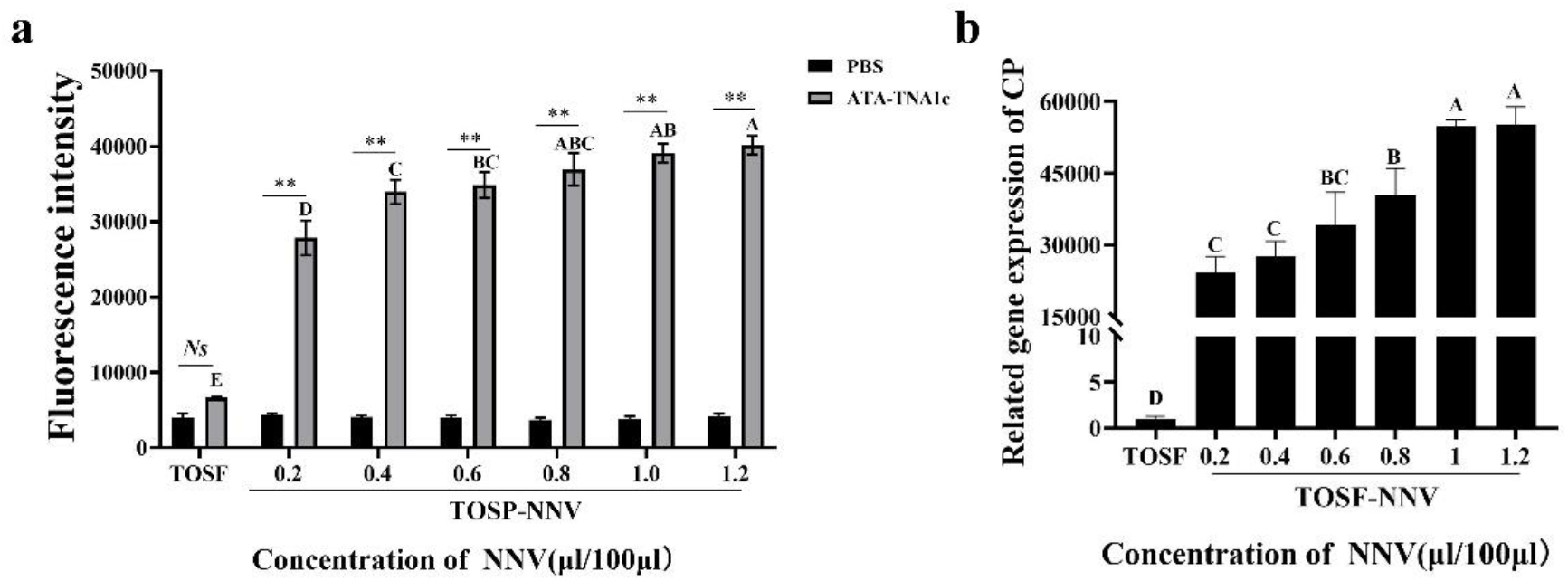
Effects of different concentrations of NNV. a. Fluorescence of TAA-TNA1c in cells infected with varying concentrations of NNV. b. qPCR analysis of CP expression levels of cells infected with different concentrations of NNV.

### Effects of different infection times of NNV

As the duration of NNV infection was increased, the effects of TOSP became more pronounced. FIG 4a shows changes to the fluorescence of TAA-TNA1c in TOSP cells infected with NNV. The fluorescence values of the probe-treated groups gradually increased with the infection time, with significant differences between the probe-treated and untreated groups at 6 to 48 h. However, at 0 h, there were no significant differences between the probe-treated and untreated groups.

**FIG 4.**
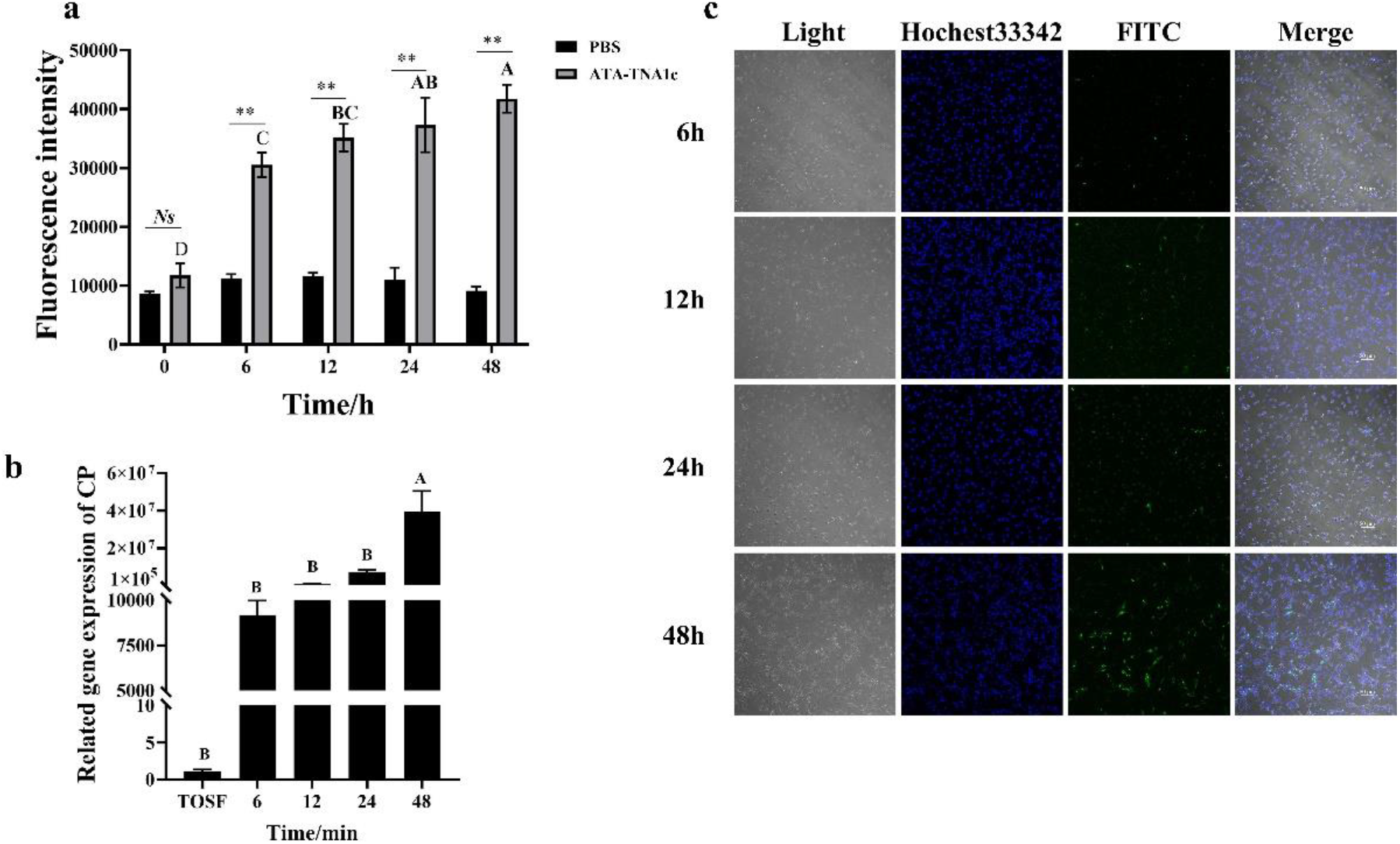
Effects of NNV infection on TOSP cells at different time points. a. Changes to fluorescence values at various time points after NNV infection. b. qPCR analysis of CP expression levels in host cells infected with NNV for different durations. c. Laser confocal microscopy assessment of fluorescence changes in host cells infected with NNV at different time intervals.

### Safe working concentrations of medicinal plant extracts

Morphological changes were observed in the TOSP cells incubated for 48 h with medicinal plant extracts or bioactive compounds. Normal TOSP cells were used as the control group, which were incubated with DMSO diluted in L15 medium to maintain normal cell growth. The viability of TOSP cells that were incubated with herbs at different concentrations was measured using the CCK-8 assay (FIG S1). The cell viability in all drug-treated groups was over 90%, and light microscopy observation (FIG S2) revealed intact and well-arranged cells with no apparent morphological differences compared to the control group. Therefore, the safe concentrations for each compound were established and are presented in TABLE 1.

**TABLE 1.** Determining the working concentrations of Herbs.

| Name | Safe concentration (µg/mL) | Name | Safe concentration (µg/mL) |
| --- | --- | --- | --- |
| SOB | 12.5 | ART | 250 |
| APR | 125 | GOI | 25 |
| XST | 12.5 | LAC | 15.6 |
| PAM | 6.25 | L-MA | 62.5 |
| WVI | 50 |  |  |

### Antiviral activity analysis of bioactive compounds by TNA1c-ATA

In this study, TAA-TNA1c was utilized to screen antiviral compounds from 9 bioactive compounds against NNV. As shown in FIG 5, five drugs exhibited antiviral effects: SOB, PAM, WVI, GOI, and L-MA (marked with “√”) . Among these, SOB, PAM, and WVI show good antiviral activity at both the safe and half-safe concentrations. However, GOI and L-MA are only effective at the safe concentration.

**FIG 5.**
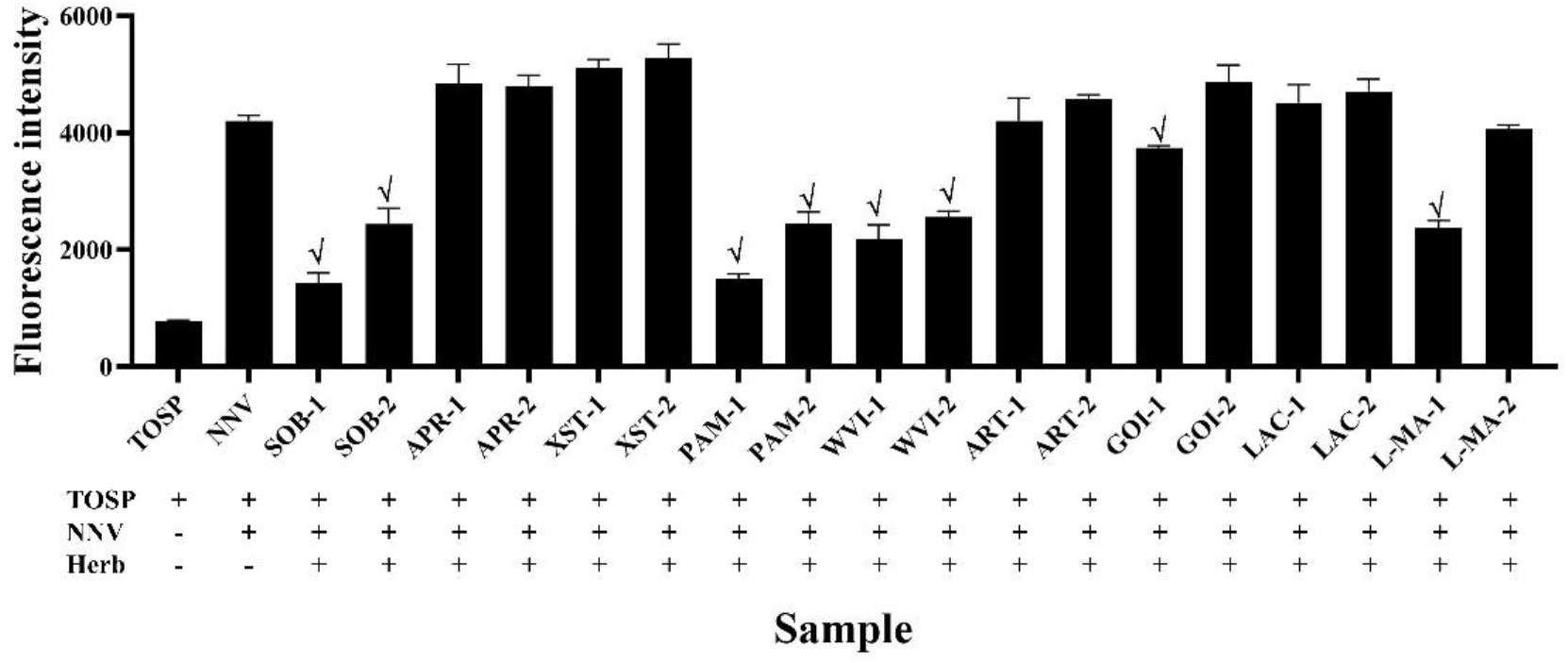
Analysis of TAA-TNA1c for high-throughput screening of anti-NNV drugs. 1 represents the drug at the safe concentration, and 2 represents the drug at half the safe concentration. Hereafter the same.

### RT-qPCR screening of the antiviral activity of medicinal plant extracts

RT-qPCR was employed to screen bioactive compounds against NNV. As shown in FIG 6, there are also five drugs that demonstrated antiviral effects: SOB, PAM, WVI, GOI, and L-MA (marked with “√”). The trend of their antiviral activity was consistent with that of the TAA-TNA1c assay results.

**FIG 6.**
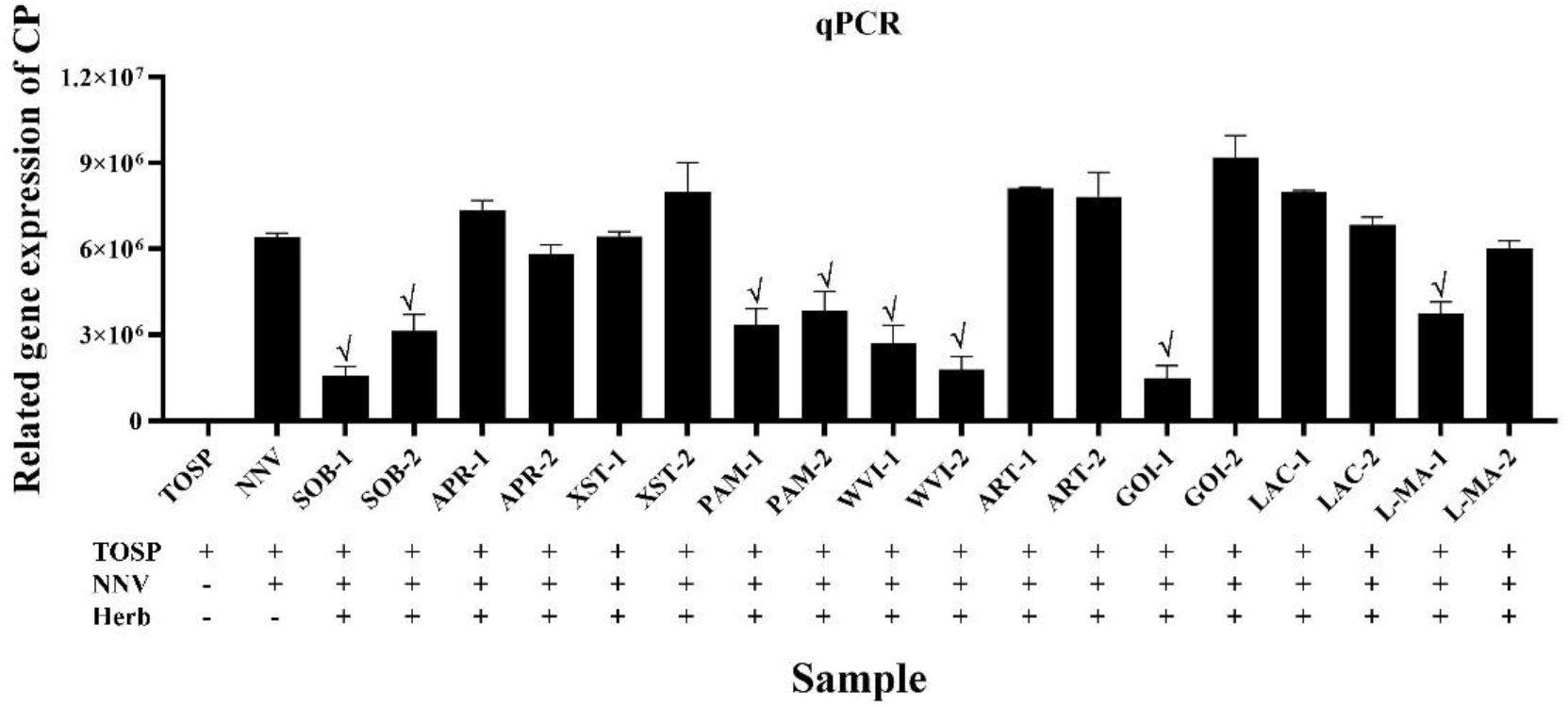
RT-qPCR analysis of the anti-NNV virus effect of medicinal plant extracts.

### Comparison of inhibition rates by TNA1c-ATA and RT-qPCR

FIG 7 illustrates the inhibition rates of various bioactive compounds against NNV infection by RT-qPCR and AHTS. As shown in FIG 7a (flow cytometry) and 7b (RT-qPCR), all effective antiviral drugs exhibited varying degrees of inhibitory activity against NNV infection, with inhibition rates ranging from approximately 35% to 77%. Specifically, samples SOB, PAM, and WVI were identified as effective antiviral agents by both methods. The discrepancy in the inhibition rate observed for sample GOI between the two detection techniques may be attributed to their distinct molecular targets. Nevertheless, this variation falls within the acceptable margin of methodological error and does not alter the overall ranking of the sample’s efficacy.

**FIG 7.**
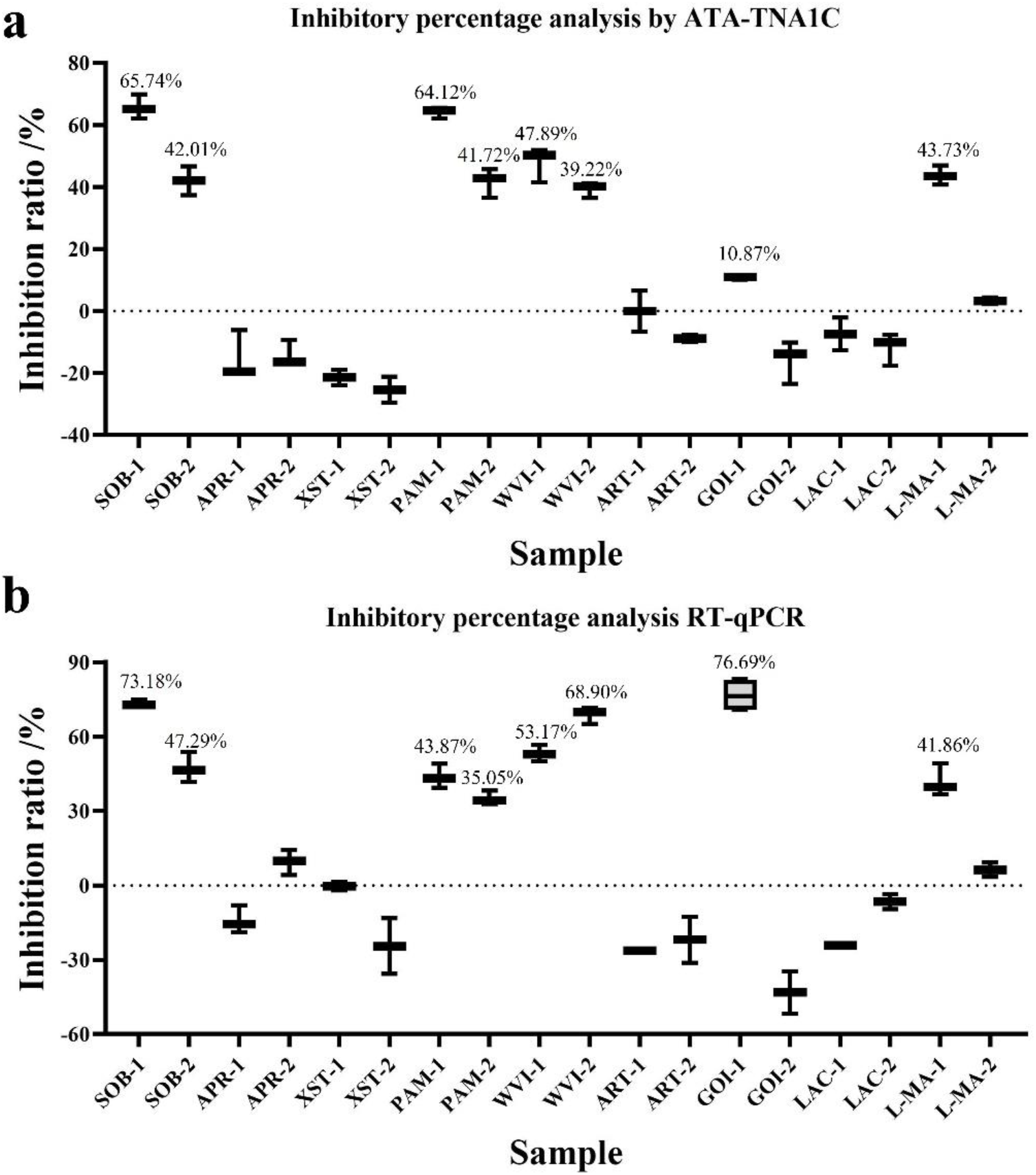
Analysis of inhibition rate of NNV infection by bioactive compounds. a. TAA-TNA1c; b. RT-qPCR.

### The method for screening anti-NNV drugs

The method for screening anti-NNV drugs is illustrated in the FIG 8. As shown, the screening of antiviral agents requires prior determination of both the safe concentration of the drug and the optimal infection concentration of NNV. After co-treating cells with NNV and the drug candidates, the antiviral effects can be evaluated using either RT-qPCR or aptamer. Among these, RT-qPCR is currently the most widely used and accurate identification method. In this study, we established a novel screening method based on aptamer probes. This approach requires only 1–2 h for the entire screening process, significantly less than the 4–5 h needed for RT-qPCR. As shown in FIG 7, the results obtained from both screening methods exhibit consistent trends, demonstrating that the high-throughput screening method developed in this study is theoretically feasible.

**FIG 8.**
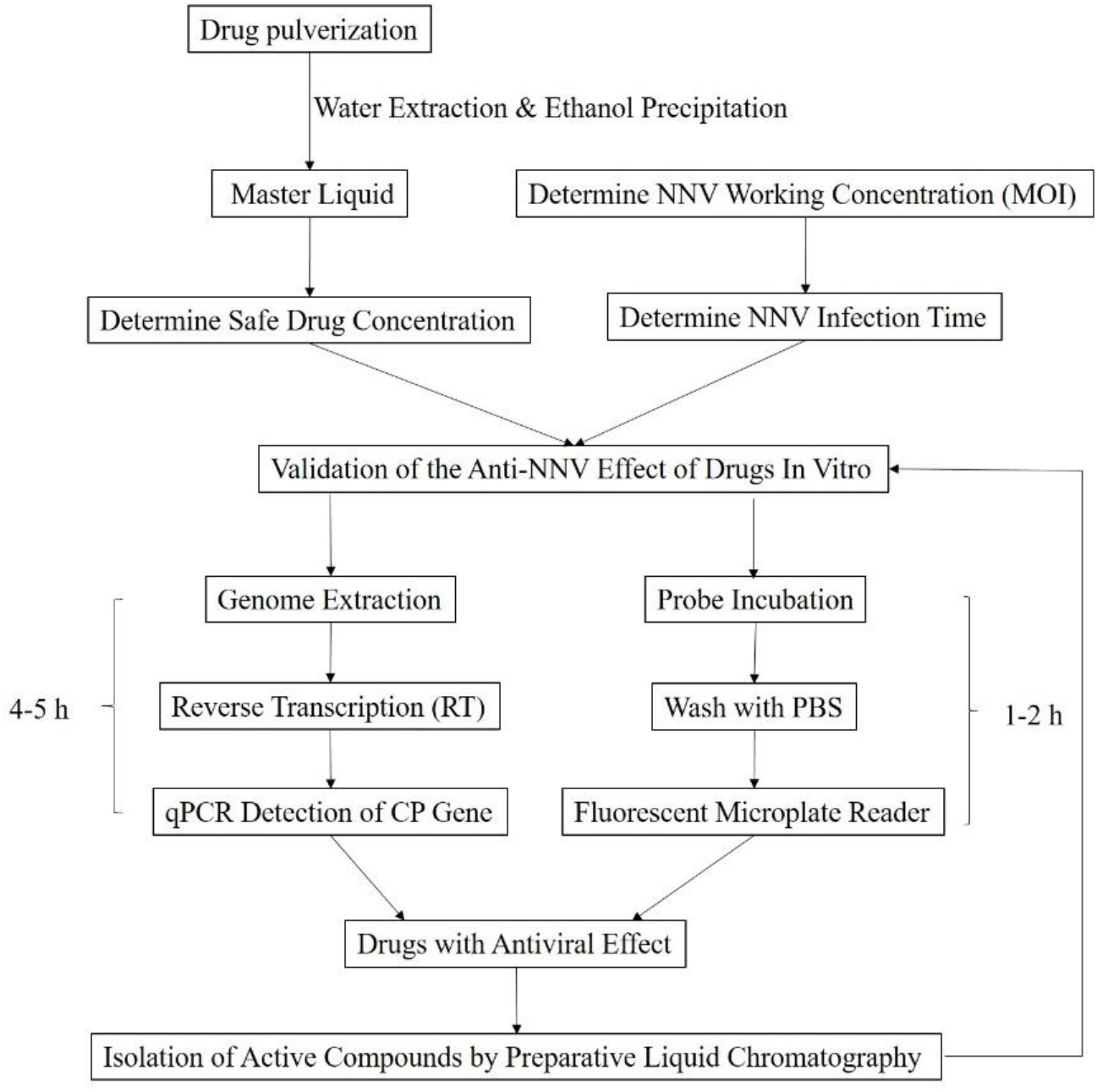
Flowchart of anti-NNV drug screening.

## DISCUSSION

NNV infection has resulted in significant economic losses to the aquaculture industry. Notably, NNV infection with high mortality rates has been reported in more than 50 wild and farmed fish species since 1986(18). These losses not only result in fish mortality but also increase operational costs, breeding time, and feed waste. An increasing number of plant-derived bioactive compounds have demonstrated antiviral activities. Therefore, screening these compounds for the development of NNV-targeting drugs is a viable, rapid, and effective approach. Currently, the most commonly used drug screening methods include RT-qPCR, cell viability assays (ATPase, kinase, and cytotoxicity) (19-21), and aptamer-based high-throughput screening methods targeting SGIV and NNV (22-24), although these methods are still considered adjunctive to RT-qPCR (3, 7).

This activatable detection method was recently introduced that maintains a low or non-fluorescent initial state, reducing background fluorescence signals, as a suitable method for fluorescence imaging of tumors(25). This activatable probe not only shortens the detection time but also enhances sensitivity. Based on previous high-throughput aptamer detection techniques, the detection aptamer was modified into an activatable probe to minimize nonspecific fluorescence to improve the accuracy of high-throughput detection (26, 27). The aptamer sgc8 has been shown to specifically recognize human acute lymphoblastic leukemia cells when modified at both ends to create an activatable probe, while retaining specificity. In this study, the TNA1c aptamer specifically recognized NNV after modification.

The detection results of aptamers are closely related to viral load, which is positively correlated with both the duration of viral infection and concentration of the virus (23, 24). In the early stages of viral infection, the virus may spread rapidly and trigger an acute immune response. Conversely, during the later stages of infection, the persistent presence of the virus leads to adaptive immune evasion in host cells, resulting in more prolonged and difficult-to-clear infections (28, 29). Furthermore, a greater initial concentration of the virus typically results in faster replication and dissemination within host cells, leading to higher viral loads and more severe clinical symptoms (23). The high-throughput detection technology developed using TAA-TNA1c in this study demonstrated time- and concentration-dependent responses when detecting NNV at varying infection concentrations and durations.

Bioactive compounds of herbs used in TCM and have been widely utilized as antiviral, antibacterial, and insecticidal agents. Various medicinal plants used in TCM, such as *Gardenia jasminoides* Ellis, *Citrus aurantium* L. (30),*Viola philippica* Cav. (31),*Arnebia euchroma, Thlaspi arvense* L.(32),*Lonicera japonica* Thunb. (6), have shown notable antiviral effects. In the present study, high-throughput detection technology with the TAA-TNA1c probe was employed to screen bioactive compounds against NNV infection. These 9 drugs were randomly selected for screening studies to evaluate their anti-NNV efficacy. The findings indicated that the detection results of TAA-TNA1c were consistent with the CP expression levels measured by RT-qPCR.

However, the high-throughput detection method significantly reduced the detection time to less than 2 h and simplified the sample extraction procedure, suggesting potential applications in drug screening.

In summary, this study optimized a critical step in antiviral drug screening by establishing a novel high-throughput method based on activatable probes. This approach reduces the entire screening process to within 2 h and yields results consistent with those obtained by RT-qPCR, demonstrating its feasibility. The proposed method minimizes operational time, reduces reagent costs, and enables high-throughput screening of a large number of compounds. Therefore, the high-throughput drug screening system holds great promise to enhance the efficiency of drug discovery.

## MATERIALS AND METHODS

### Cells, viruses and reagents

Splenic fibroblasts of the golden pompano (*Trachinotus ovatus*) (TOSP cell line), established by our laboratory (33), were cultured and propagated in Leibovitz’s L15 medium (Gibco™; Thermo Fisher Scientific, Waltham, MA, USA) supplemented with 10 % fetal bovine serum (Gibco™) at 28°C. The virulence of NNV isolated from *T. ovatus*(34) and SGIV, also isolated by our laboratory, was verified by regression infection *in vivo*. Both cells and viruses were stored in a pathogen repository at -80°C in the Key Laboratory of Aquatic Biotechnology and Modern Ecological Aquaculture, Guangxi Academy of Sciences (Nanning, China).

*Syringa oblate* (SOB), *Astragalus propinquus* (APR), *Xanthium strumarium* (XST), *Phellodendri amurensis cortex* (PAM), *Wurfbainia villosa* (WVI), Artemisinin (ART), Garlic oil (GOI), Lipoic acid (LAC), L-Malic acid (L-MA) were purchased from Shanghai Macklin Biochemical Technology Co., Ltd.

### Preparation of probes and primers

The aptamer TNA1c, which targets cells infected by NNV isolated from *T. ovatus*, was obtained by SELEX (Systematic Evolution of Ligands with EXponential enrichment) technology, as described in a previous study (35). The sequence of the activatable aptamer TNA1c (TAA-TNA1c) is 5′-FAM-CCA ACC TTT TTT TTT TTT TTT TTT TGC TCG GGG CTC GAT ATT GTA AAG GGA GTG TGT TTA GGA GGA CGT GGT TGG-BHQ1-3′. The primers used in the experiments are listed in TABLE 2. All sequences and primers were synthesized by Beijing AuGCT DNA-SYN Biotechnology Co., Ltd. (Beijing, China). Prior to experimentation, the aptamers were subjected to a heat denaturation step at 92°C for 5 min, then cooled in ice bath for 5 min and stored on ice.

**TABLE 2.**
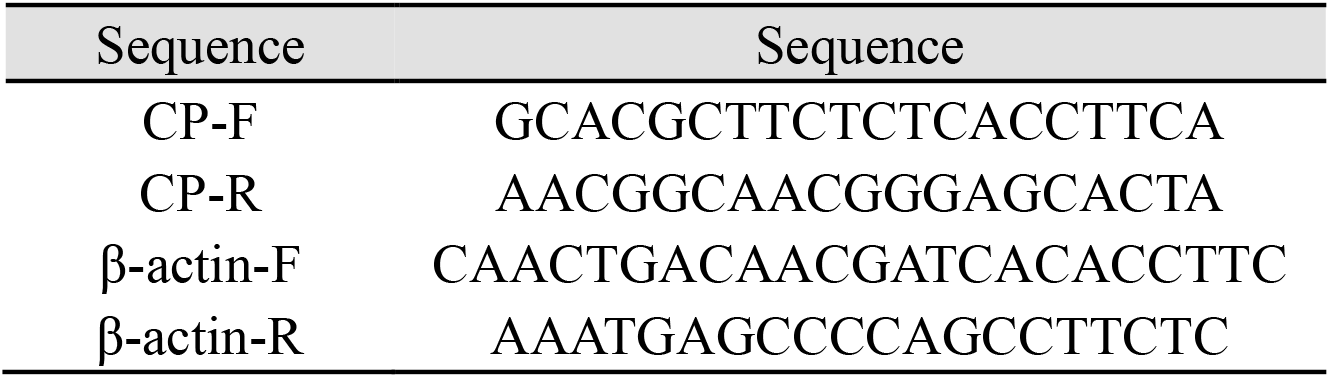
Primers for testing.

### Laser Confocal Analysis of TAA-TNA1c Specificity

TOSP cells (1 × 10^5^/mL) were seeded on a 35-mm glass-bottom dish (Cellvis; catalog number D35-14-1-N) and cultured at 28°C for 18 h. Afterward, the cells were cultured at 28°C for 48 h with NNV (multiplicity of infection [MOI] = 1), as the experimental group, SGIV (MOI = 1), as a negative control group, or left untreated, as a control group. The denatured TAA-TNA1c probes were added to each group and incubated on ice for 30 min. Then, Hoechst 33342, a fluorescent dye for DNA staining, was added to each group. After 15 min, fluorescence was measured at 490/525 nm with a laser confocal microscope.

### Determination of NNV Infection Concentration

TOSP cells (1×10^6^/mL) were seeded in the wells of 24-well plate and cultured at 28°C for 18 h. Afterward, the cells were infected with NNV (0.2, 0.4, 0.6, 0.8, 1.0, and 1.2 μL/100 μL) or cultured in serum-free L15 medium as a control group. After 48 h of infection, tissue RNA was extracted using TRIzol reagent and reverse transcribed into cDNA. Then, expression of the CP gene was quantified by RT-qPCR.

Additionally, TOSP cells (1 × 10^6^/mL) were seeded in the wells of 96-well plate and cultured at 28°C for 18 h. Afterward, the cells were infected with NNV (0.2, 0.4, 0.6, 0.8, 1.0, and 1.2 μL/100 μL) or cultured in serum-free L15 medium as a control group. At 48 h post-infection, 10 μL of TAA-TNA1c were added to each well and incubation was continued for 30 min at 4°C. Fluorescence was measured with a microplate reader at 490/525 nm.

### Determination of NNV infection duration

TOSP cells (1 × 10^6^/mL) were seeded in the wells of a 24-well plate and cultured at 28°C for 18 h. Then, the cells were infected with NNV diluted to 0.2 μL/100 μL for 6, 12, 24, and 48 h, or cultured in serum-free L15 medium as a control group. Afterward, total RNA was extracted as described in section 2.4.

In addition, TOSP cells (1 × 10^6^/mL) were seeded in the wells of a 96-well plate and cultured at 28°C for 18 h. Then, the cells were infected as described in section 2.4.

### Analysis of the safe working concentrations of medicinal plant extracts

The effects of different concentrations of medicinal plant extracts on cell viability were determined as described previously (36). Briefly, TOSP cells were seeded in a 96-well plate (Corning) at a density of 1 × 106 cells per well and cultured for 18 h at 28 °C. Subsequently, the cells were treated with different concentrations of the extracts and incubated for 48 h at 28 °C. In the control group, GS cells were incubated with L15 medium containing diluted DMSO. Cytotoxicity was preliminarily evaluated by light microscopy at 48 h. The cells were then washed twice with phosphate-buffered saline (PBS) and incubated for 4 h at room temperature with CCK-8 solution (Beyotime, Shanghai, China) diluted to 10% in PBS. Cell viability was determined by measuring the absorbance at 450 nm using a spectrophotometer (Thermo Fisher Scientific, Waltham, MA, USA). All experiments were performed in quadruplicate and repeated three times independently. Data are presented as means ± standard deviation (SD).

### Antiviral activity analysis of bioactive compounds by TNA1c-ATA

TOSP cells (1 × 106/mL) were seeded in a 96-well plate and cultured at 28°C for 18 h. Afterward, the cells were infected with NNV (MOI = 1) and treated with medicinal plant extracts at their highest safe concentration and a half-dilution of that concentration. The cells were infected with NNV (MOI = 1) only as the positive control group. Cells cultured in serum-free L15 medium served as the control group. Each treatment group included four replicates. After 48 h, the supernatant was removed, and denatured probes (500 nM) were added to the wells for 30 min at 4°C. The wells were then gently washed once with 200 µL of phosphate-buffered saline (PBS). Subsequently, 100 µL of PBS was added to each well, and fluorescence was measured using a fluorescence microplate at excitation/emission wavelengths of 495/535 nm.

### RT-qPCR screening of antiviral activity of medicinal plant extracts

TOSP cells (1 × 106/mL) were seeded in a 12-well plate and incubated overnight. Control group received serum-free L-15 medium only, while positive control group was treated with NNV solution only. The experimental groups were treated with NNV in combination with either the highest safe concentration of medicinal plant extracts or its half-dilution, with four replicates per group. After 48 h of treatment, cells were collected and total RNA was extracted using TRIzol reagent, followed by reverse transcription into cDNA. CP gene expression was then quantified by RT-qPCR.

### Data Analysis

Data analysis was performed with IBM SPSS Statistics for Windows (version 26.0; IBM Corporation, Armonk, NY, USA) and graphs were generated with Prism 9.0 software (GraphPad Software, LLC, San Diego, CA, USA). The data among groups were compared by two-way analysis of variance. A probability (*p*) value ≤ 0.05 was considered statistically significant.

## ACKNOWLEDGMENTS

This work was financially supported by Natural Science Foundation of Guangxi (2022GXNSFBA035521 to Q.Y. and 2022JJA130281 to Q.Y.), National Natural Science Foundation of China (32373175 to P.L.), Guangxi Innovation Team Project of National Modern Agricultural Industrial Technology System (nycytxgxcxtd-2021-08-02 to P.L.), Guangxi Science and Technology Program (GUIKEFA[2024]102-2 to P.L.), Laibin Science and Technology Program (Laikechan 241519 to D.W.).

P.L. conducted this work. Y.G. and W.X. performed the experiment. J.G., L.H., H.C., Y.Z., J.C., X.Z., S.H. and J.X. analyzed data. Z.L. and Q.Y. wrote the paper.

## FUNDING

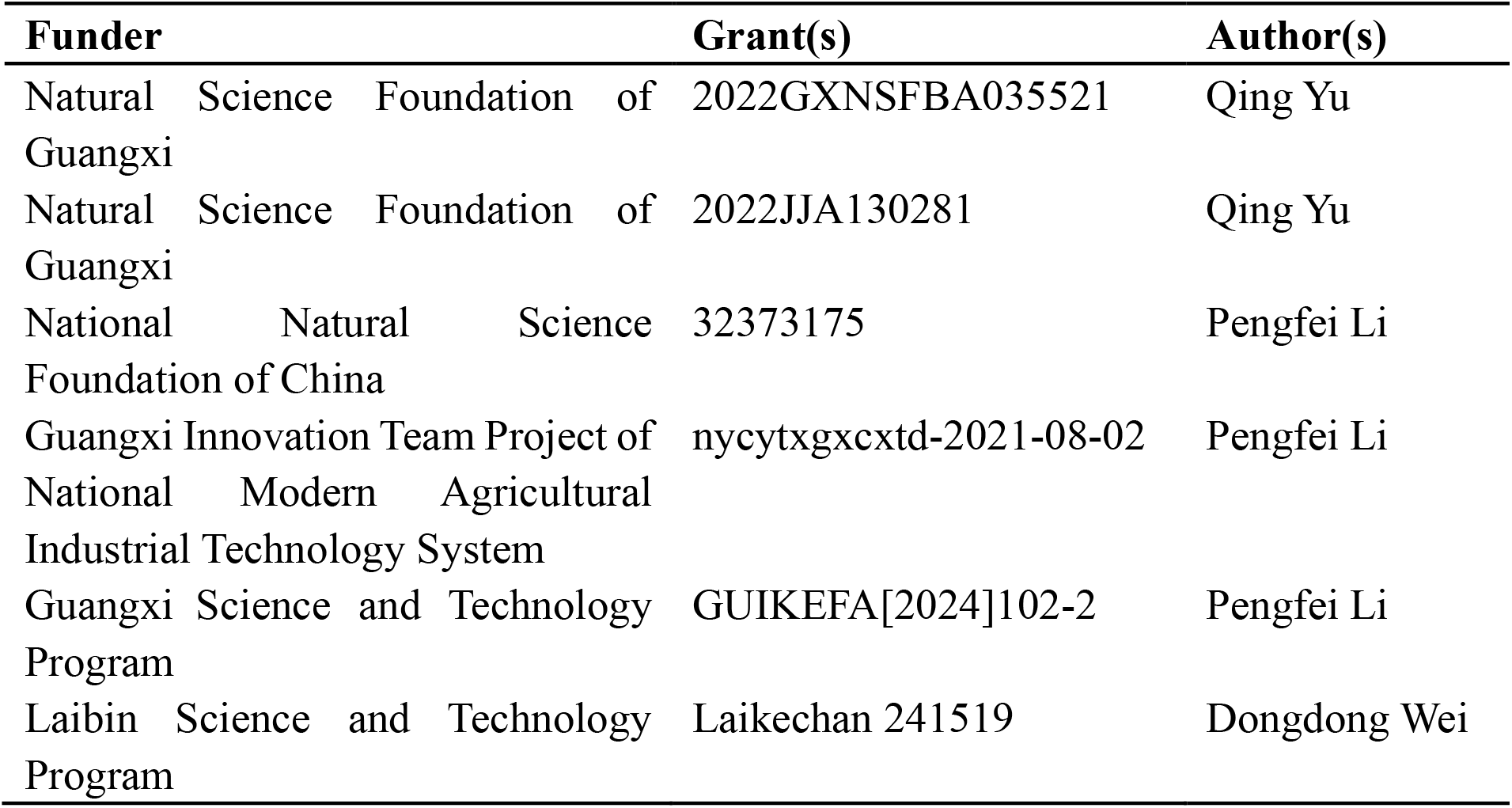

## REFERENCE

1. Huang L, Liu MZ, Yu Q, Han SY, Wei DD, Shi JG, Wei HL, Li PF. 2025. Study and analysis of pathogen co-infection in grouper culture. Journal of fisheries of China 49:039416.

2. Li JX, Li SQ, Liu ZL, Huang DC, Li X, Liang JM, Wang ZP, Teng YL, Liu P, Chai H, Huang L, Zhao MM, Li PF. 2025. Future Path of Guangxi Fishery: Green Intensive Smart Aquaculture and Deep Processing Model of Characteristic Aquatic Products. Journal of Guangxi Academy of Sciences 41:1–10.

3. Li MM, Liu MZ, Wei HL, Huang L, Yu Q, Huang SS, Li JY, Li PF. 2022. Antiviral activities of Glycyrrhiza uralensis components against Singapore grouper iridovirus. Journal of the World Aquaculture Society 53:894–909.

4. Liu MZ, Xiao HH, Zhang Q, Wu ST, Putra DF, Xiong XY, Xu MZ, Dong LF, Li SQ, Yu Q, Li PF. 2019. Antiviral abilities of Curcuma kwangsiensis ingredients against grouper iridoviral infection in vitro and in vivo. Aquaculture Research 51:351–361.

5. Liu MZ, Yu Q, Xiao HH, Yi Y, Cheng H, Putra DF, Huang YM, Zhang Q, Li PF. 2020. Antiviral activity of Illicium verum Hook. f. extracts against grouper iridovirus infection. Journal of Fish Diseases 43:531–540.

6. Liu MZ, Yu Q, Yi Y, Xiao HH, Putra DF, Ke K, Zhang Q, Li PF. 2020. Antiviral activities of Lonicera japonica Thunb. Components against grouper iridovirus in vitro and in vivo. Aquaculture 519:734882.

7. Cheng Y, Liu MZ, Yu Q, Huang SS, Han SY, Shi JG, Wei HL, Zou JW, Li PF. 2023. Effect of EGCG Extracted from Green Tea against Largemouth Bass Virus Infection. Viruses 15:151.

8. Jiang HF, Chen C, Jiang XY, Shen JL, Ling F, Li PF, Wang GX. 2022. Luteolin in Lonicera japonica inhibits the proliferation of white spot syndrome virus in the crayfish Procambarus clarkii. Aquaculture 550:737852.

9. Huang L, Li MM, Wei HL, Yu Q, Huang SS, Wang TX, Liu MZ, Li PF. 2022. Research on the indirect antiviral function of medicinal plant ingredient quercetin against grouper iridovirus infection. Fish and Shellfish Immunology 124:372–379.

10. Song Y, Zhang Y, Xiao S, Li P, Lu L, Wang H. 2024. Akt inhibitors prevent CyHV-2 infection in vitro. Fish and Shellfish Immunology 154:109940.

11. Yu Q, Liu MZ, Xiao HH, Yi Y, Cheng H, Putra DF, Li SQ, Li PF. 2020. Selection and Characterization of Aptamers for Specific Detection of Iridovirus Disease in Cultured Hybrid Grouper (Epinephelus Fuscoguttatus♀ × E. Lanceolatus♂). Chinese Journal of Analytical Chemistry 48:650–661.

12. Zhou LL, Li PF, Yang M, Yu YP, Huang YH, Wei JG, Wei SN, Qin QW. 2016. Generation and characterization of novel DNA aptamers against coat protein of grouper nervous necrosis virus (GNNV) with antiviral activities and delivery potential in grouper cells. Antiviral Research 129:104–114.

13. Zhou LL, Wang SW, Yu Q, Wei SN, Liu MZ, Wei JG, Huang YH, Huang XH, Li PF, Qin QW. 2020. Characterization of Novel Aptamers Specifically Directed to Red-Spotted Grouper Nervous Necrosis Virus (RGNNV)-Infected Cells for Mediating Targeted siRNA Delivery. Front Microbiology 11:660.

14. Zhang XY, Zhang ZM, Li JR, Huang XH, Wei JG, Yang JH, Guan LF, Wen XZ, Wang SW, Qin QW. 2022. A Novel Sandwich ELASA Based on Aptamer for Detection of Largemouth Bass Virus (LMBV). Viruses 14:945.

15. Lee S, Xie J, Chen XY. 2010. Activatable Molecular Probes for Cancer Imaging. Current Topics in Medicinal Chemistry 10: 1135–1144.

16. Shi H, He XX, Wang K, Wu X, Ye XS, Guo QP, Tan WH, Qing ZH, Yang XH, Zhou B. 2011. Activatable aptamer probe for contrast-enhanced in vivo cancer imaging based on cell membrane protein-triggered conformation alteration. Proceedings of the National Academy of Sciences 108:3900–5.

17. Zeng ZH, Tung CH, Zu YL. 2014. A cancer cell-activatable aptamer-reporter system for one-step assay of circulating tumor cells. Molecular Therapy Nucleic Acid 3:e184.

18. Mohammad JZ. 2020. Viral nervous necrosis disease, p 673–703, Emerging and reemerging viral pathogens doi:10.1016/B978-0-12-819400-3.00030-2. Academic Press.

19. Severson WE, McDowell M, Ananthan S, Chung DH, Rasmussen L, Sosa MI, White EL, Noah J, Jonsson CB. 2008. High-throughput screening of a 100,000-compound library for inhibitors of influenza A virus (H3N2). Journal of biomolecular screening 13:879–87.

20. Gong E, Ivens T, Van den Eynde C, Hallenberger S, Hertogs K. 2008. Development of robust antiviral assays for profiling compounds against a panel of positive-strand RNA viruses using ATP/luminescence readout. Journal of Virological Methods 151:121–5.

21. Li QJ, Maddox C, Rasmussen L, Hobrath JV, White LE. 2009. Assay development and high-throughput antiviral drug screening against Bluetongue virus. Antiviral Research 83:267–73.

22. Liu MZ, Xiao HH, Wu ST, Yu Q, Li PF. 2020. Aptamer-based high-throughput screening model for medicinal plant drugs against SGIV. Journal of Fish Diseases 43:1479–1482.

23. Wei HL, Guo ZB, Long Y, Liu MZ, Xiao J, Huang L, Yu Q, Li PF. 2022. Aptamer-Based High-Throughput Screening Model for Efficient Selection and Evaluation of Natural Ingredients against SGIV Infection. Viruses 14:1242.

24. Wei HL, Liu MZ, Ke K, Xiao SY, Huang L, He QY, Mo CP, Pang H, Xiao GZ, Li PF, Yu Q. 2022. Study on aptamer based high throughput approach identifies natural ingredients against RGNNV. Journal of Fish Diseases 45:1711–1719.

25. Deng JQ, Tian F, Liu C, Liu Y, Zhao S, Fu T, Sun JS, Tan WH. 2021. Rapid One-Step Detection of Viral Particles Using an Aptamer-Based Thermophoretic Assay. Journal of the American Chemical Society 143:7261–7266.

26. Zhao BJ, Wu P, Zhang H, Cai CX. 2015. Designing activatable aptamer probes for simultaneous detection of multiple tumor-related proteins in living cancer cells. Biosensors and Bioelectronics 68:763–770.

27. Lai ZQ, Tan JT, Wan RR, Tan J, Zhang ZH, Hu ZX, Li JP, Yang W, Wang YW, Jiang YF, He J, Yang N, Lu XL, Zhao YX. 2017. An ‘activatable’ aptamer-based fluorescence probe for the detection of HepG2 cells. Oncology Reports 37:2688–2694.

28. Huang YH, Huang XH, Wang SW, Yu YP, Ni SW, Qin QW. 2018. Soft-shelled turtle iridovirus enters cells via cholesterol-dependent, clathrin-mediated endocytosis as well as macropinocytosis. Archives of Virology 163:3023–3033.

29. Huang XH, Yu T, Zhao Y, Huang YH, Qin QW. 2024. Research progress on the regulation and utilization of host cell metabolism by aquatic animal viruses. Acta Hydrobiologica Sinica 49:1–23.

30. Yang Y, Cheng H, Yan H, Wang PZ, Rong R, Zhang YY, Zhang CB, D. RK, Rong LJ. 2017. A cell-based high-throughput protocol to screen entry inhibitors of highly pathogenic viruses with Traditional Chinese Medicines. Journal of Medical Virology 89:908–916.

31. Yu Q, Liu MZ, Xiao HH, Wu ST, Qin XL, Lu ZJ, Shi DQ, Li SQ, Mi HZ, Wang YB, Su HF, Wang TX, Li PF. 2019. The inhibitory activities and antiviral mechanism of Viola philippica aqueous extracts against grouper iridovirus infection in vitro and in vivo. Journal of Fish Diseases 42:859–868.

32. Ho TY, Wu SL, Lai IL, Cheng KS, Kao ST, Hsiang CY. 2003. An in vitro system combined with an in-house quantitation assay for screening hepatitis C virus inhibitors. Antiviral Research 58:199–208.

33. Huang L, Liu MZ, Lu XH, Kuang JH, Zhu LB, Wei YY, Ke K, Wang H, Lv TY, Xiao J, Yu Q, Li PF. 2024. Establishment a cell line of splenic fibroblasts from golden pompano and its application in heavy metal toxicology and pathogen infection. Aquaculture and Fisheries doi:10.1016/j.aaf.2024.09.001.

34. Li PF, Yu Q, Li F, Qin XL, Dong DX, Chen B, Qin QW. 2018. First identification of the nervous necrosis virus isolated from cultured golden pompano (Trachinotus ovatus) in Guangxi, China. Journal of Fish Diseases 41:1177–1180.

35. Yu Q, Liu MZ, Wei SN, Wu ST, Xiao HH, Qin XL, Su HF, Li PF. 2019. Characterization of ssDNA aptamers specifically directed against Trachinotus ovatus NNV (GTONNV)-infected cells with antiviral activities. Journal of General Virology 100:380–391.

36. Xu WQ, Yu JY, Huang L, Yu Q, Liu MZ, Wang SW, Sun Y, Han SY, Qin QW, Li PF. 2025. Action mechanism of Semen Ziziphi Spinosae alcohol extracts and betulinic acid against nervous necrosis virus. Journal of Southern Agriculture 56:1787–1801.

